# BacNeMu: neutral mutation spectra reconstruction pipeline for bacteria

**DOI:** 10.64898/2026.06.30.735404

**Authors:** Alexey Skudnov, Eldar Badamshin, Bogdan Efimenko, Konstantin Popadin, Konstantin Gunbin, Stepan Denisov

**Affiliations:** Center for Mitochondrial Functional Genomics, Immanuel Kant Baltic Federal University, Kaliningrad, Russia; Limnological Institute, Siberian Branch of the Russian Academy of Sciences, Irkutsk, Russia; CellKinetica, Switzerland; The University of Manchester, Manchester, UK

## Abstract

The mutational spectrum is an increasingly important molecular phenotype that quantitatively describes mutagenesis in a given gene and species, enabling future comparative analyses to reveal differences in underlying mutagenic processes, whether internal, such as DNA repair processes, or external, such as ecological niches and conditions. Mutation accumulation experiments, although time-consuming and costly, remain the standard approach for reconstructing bacterial neutral mutation spectra. Here, we present BacNeMu, a phylogenetically informed pipeline that reconstructs neutral mutational spectra of bacterial genomes using open databases GTDB^1^, AnnoTree^2^ and KEGG Orthology^3^, building on previously developed NeMu pipeline^4^. BacNeMu reconstructs mutation spectra that closely match mutation accumulation experiments results while requiring substantially less time, enabling comparative analyses across diverse bacterial taxa. Applied to obligate aerobes and anaerobes, BacNeMu recovered the expected excess of T:A>C:G transitions, consistent with oxidative-damage-associated mutational patterns previously described in mitochondrial genomes and yeast single-strand^5,6^. We further asked if any other ecologic factors influence a mutational spectrum. As a pilot we compared three species living under different temperatures: one strong thermophile - *Thermotoga maritima*^7^, one psychrophile - *Clostridium algidicarnis*^8^, and one with intermediate temperature tolerance - *Psychrobacter sanguinis*^9^. In the thermophile, the relative frequency of T:A>C:G substitutions was higher than in the psychrophile, although C:G>T:A transitions predominate across all three species. BacNeMu provides a rapid, phylogenetically informed framework for generating biologically meaningful mutation spectra from open databases.

**Graphical abstract:** 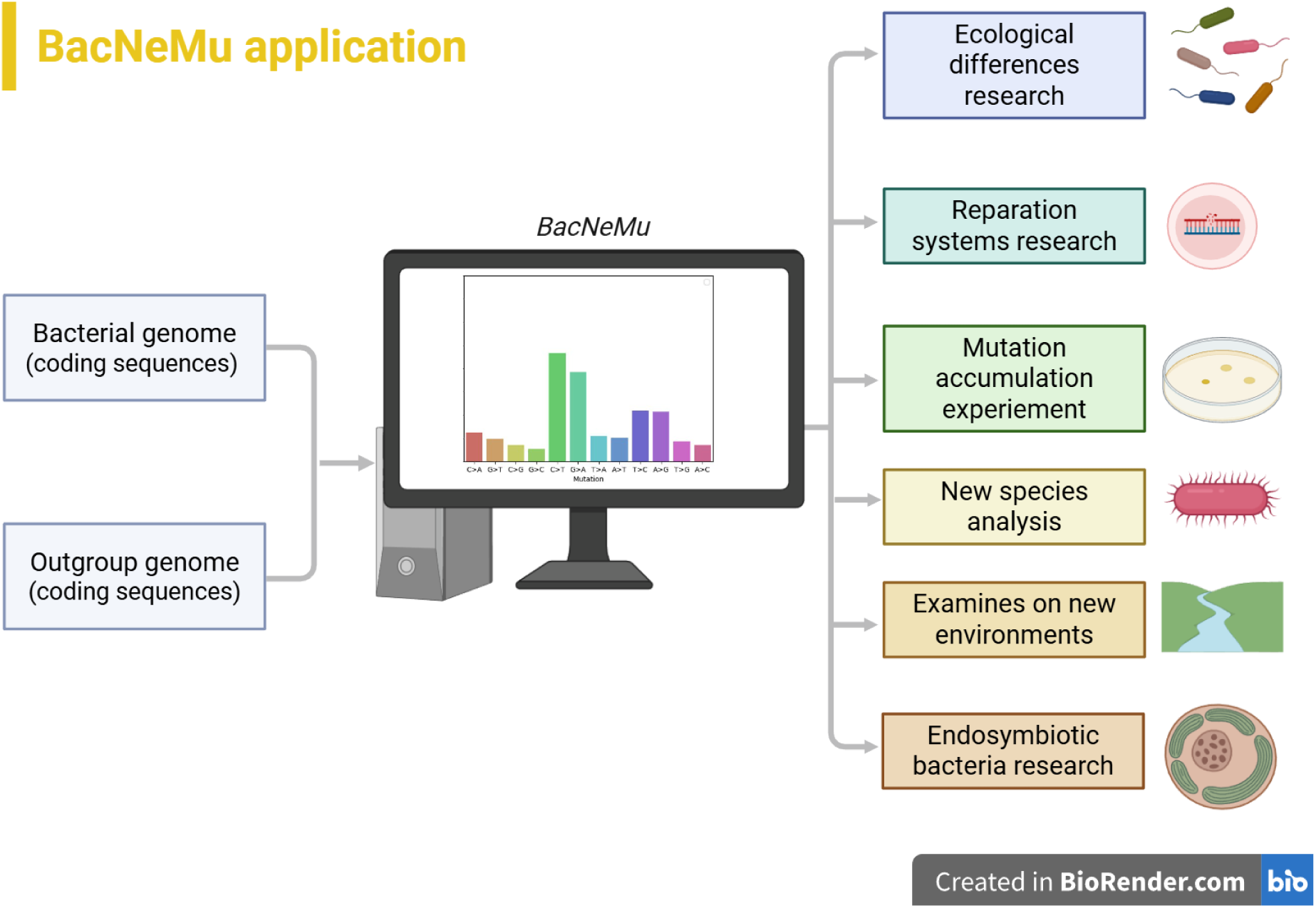

## Introduction

A neutral mutational spectrum quantifies the relative frequencies of nucleotide substitutions that accumulate in a genome under conditions where natural selection is minimised. Under such regimes, two evolutionary forces predominate: mutagenesis — the biochemical generation of sequence alterations — and genetic drift — the stochastic fluctuation of allele frequencies across generations. Neutral spectra are informative because they capture the integrated effects of endogenous DNA maintenance processes, including replication, recombination, and repair, together with exogenous ecological pressures that compromise DNA integrity^10^. The principal methodological challenge lies in identifying genomic positions that evolve in an effectively neutral manner. The NeMu pipeline addresses this by targeting third-codon positions of four-fold degenerate codons: sites at which any nucleotide substitution leaves the encoded amino acid unchanged, rendering such substitutions largely invisible to purifying selection^4^. Because these sites freely accumulate substitutions, they preserve a near-neutral record of the mutational spectrum along a phylogeny — a signal that can be read backwards to infer ancestral mutational processes. This principle — that mutational spectra encode information about their source — finds precedents outside bacterial genomics. In oncogenetics, characteristic mutational signatures now enable the classification of tumour types directly from sequencing data^11,12^, while early analyses of mitochondrial DNA demonstrated that an elevated rate of adenine-to-guanine (A>G) transitions is associated with oxidative stress and, over longer timescales, with organismal ageing^5^. NeMu extends this logic to the reconstruction of bacterial mutational histories from phylogeny.

In bacteria, the relationship between genomic nucleotide composition and the underlying mutational spectrum remains incompletely resolved^10,13^. Two well-documented yet puzzling observations provide the motivation for the present work. The first concerns oxygen availability: obligate aerobic bacteria tend to possess GC-rich genomes^14^, whereas obligate anaerobes are predominantly AT-rich^15^. However, the canonical oxygen-induced lesion, G→T transversion^14^, should erode GC content — yet obligate aerobes, which face chronic oxidative stress, remain GC-richer than anaerobes. The forces reconciling this contradiction are unknown. The second concerns growth temperature: thermophilic bacteria consistently exhibit elevated genomic GC content relative to mesophiles and psychrophiles, leading to the hypothesis that GC-rich DNA is selectively favoured at high temperatures owing to its greater thermal stability^16^. For both patterns, the same causal question is debated: do these compositional biases reflect environment-specific selective pressures on nucleotide composition, or do they arise from systematic biases in the mutational process itself — such that organisms inhabiting distinct oxygen or temperature regimes experience qualitatively different spectra of spontaneous mutations^17,18^. Resolving this question demands a method capable of rapidly and reliably reconstructing neutral mutational spectra across a broad and phylogenetically diverse set of bacterial species, thereby enabling the mutational baseline to be disentangled from the footprint of natural selection.

Comparative analyses of mutational spectra across species must contend with a well-recognised complication: closely related species often resemble one another in genomic traits not because they adapted independently to similar ecological conditions, but because they inherited those traits from a recent common ancestor — a phenomenon termed phylogenetic inertia^19^. Failure to account for this non-independence can generate spurious associations between ecological variables and mutational patterns, creating the misleading impression that a particular environmental factor drives a given mutational signature, when the underlying signal may instead reflect shared ancestry within a single clade^20,21^. A rigorous method for reconstructing neutral mutation spectra across diverse taxa should therefore either ensure that the sampled species are widely dispersed across the bacterial tree of life — so that no single clade dominates any ecological category — or incorporate statistical frameworks, such as phylogenetic generalised least squares, that explicitly correct for the covariance introduced by shared evolutionary history^22^.

To address these challenges, we introduce BacNeMu — a bioinformatic pipeline designed for the rapid and accurate reconstruction of neutral mutational spectra in bacteria. BacNeMu extends the NeMu framework^4^ from eukaryotes to bacteria by addressing a fundamental obstacle: unlike eukaryotic genomes, bacterial genomes are shaped by horizontal gene transfer (HGT)^23^ and gene duplication events^24^. Both processes generate paralogous sequences — genes that share similarity not because they diverged through vertical speciation, but because they were acquired from an unrelated donor (HGT) or copied within the same genome (duplication). When such paralogs are inadvertently included in a multiple-sequence alignment, the reconstructed gene tree conflates vertical and non-vertical signals: internal nodes no longer correspond to speciation events, and synonymous substitutions are mapped to incorrect branches, biasing the inferred mutational spectrum. Robust reconstruction therefore requires that the analysis be restricted to one-to-one (1:1) orthologs — homologous genes whose divergence is attributable to speciation alone and whose gene tree faithfully mirrors the organismal phylogeny.

BacNeMu operates in two modes. In fully automatic mode, the user provides only a list of species names in GTDB format^1^; the pipeline then retrieves all annotated genomes of the corresponding strains from a pre-loaded database, identifies one-to-one orthologous gene clusters, selects a suitable outgroup strain using the AnnoTree phylogenetic framework^2^, and reconstructs a species-level six-component mutational spectrum — covering all six possible base-pair substitutions — from synonymous changes inferred along internal branches of gene trees built with IQ-TREE^25^. In manual mode, users may supply their own gene sets, with outgroup selection left to their discretion. A typical run completes within one to eight hours, depending on the number of input strains and technical parameters of the computer, making BacNeMu a practical high-throughput alternative to long-term mutation accumulation experiments.

To validate the pipeline, we compared BacNeMu-derived spectra with previously published mutational spectra obtained from mutation accumulation experiments (MAE) across six bacterial species: *Escherichia coli*, *Bacillus subtilis*, *Vibrio cholerae*, *Aliivibrio fischeri*, *Burkholderia cenocepacia* and *Mycobacterium smegmatis*. For synonymous substitutions in particular, the BacNeMu-inferred spectra showed broad agreement with the experimentally determined MAE profiles, indicating that the pipeline captures the underlying neutral mutational process with reasonable fidelity. As an additional benchmark, we applied MutTui^26^ — an alternative computational method that employs phylogenetic reconstruction to infer the direction of mutational changes from single-nucleotide polymorphisms — to the same dataset, normalising the resulting spectra against the nucleotide composition of the reference genome used by MutTui to ensure comparability. The two computational approaches yielded highly concordant spectra across all six substitution types, as assessed by cosine similarity, further confirming the robustness of phylogenetically inferred mutational profiles. As a pilot application of BacNeMu, we investigated whether obligate aerobic and obligate anaerobic bacteria differ systematically in their neutral mutational spectra, and whether growth temperature leaves a detectable imprint on the spectra of thermophiles and psychrophiles. For the aerobic–anaerobic comparison, prior work on mitochondrial DNA has linked an elevated rate of A:T>G:C transitions to oxidative stress^5^, and a similar excess was observed in the BacNeMu-inferred spectra of obligate aerobes, consistent with the hypothesis that oxygen availability shapes the mutational landscape of bacteria.

## Materials and methods

To enforce this restriction, BacNeMu implements an automated orthology-filtering step that integrates three publicly available resources: GTDB R214 for curated bacterial genome sequences and taxonomic metadata³, AnnoTree r214 for a reference prokaryotic phylogeny linked to the GTDB taxonomy⁴, and KEGG Orthology — a database of functionally annotated clusters of homologous proteins, queried here via kofam-scan v.1.3.0 (a hidden Markov model-based search tool)⁵.

The pipeline requires minimal input: the user provides a list of species of interest formatted according to the GTDB taxonomy (e.g., s Escherichia coli, where the s prefix denotes the species rank)^1^. From this list, BacNeMu queries GTDB to retrieve all available strains assigned to each species together with their unique genome identifiers (gtdb_id). For every identified strain, the pipeline downloads the full set of annotated protein-coding gene sequences (nucleotide FASTA, .fna) and the corresponding translated protein sequences (amino acid FASTA, .faa). Individual genes are subsequently matched across the two files by their internal sequence identifiers, ensuring that each nucleotide sequence is paired with its correct translation for downstream orthology assignment.

Accurate rooting is essential for inferring mutation directionality: an unrooted tree cannot distinguish an A>G substitution from its reverse complement G>A, because both are equally parsimonious without a known ancestral state at the root. BacNeMu therefore automatically selects an outgroup strain using the AnnoTree prokaryotic phylogeny^2^, which links GTDB reference genomes to their positions on a comprehensive bacterial tree of life. For each focal species, the pipeline identifies the corresponding node in AnnoTree, then — via the Python library ETE3^27^ — locates the nearest reference genome assigned to a different species, choosing the neighbour with the shortest phylogenetic distance. When multiple candidate outgroups are equally distant, one is selected at random. The nucleotide and protein sequences of the selected outgroup strain are retrieved from GTDB and merged with the focal-species sequences (Fig. 1)

**Figure 1.**
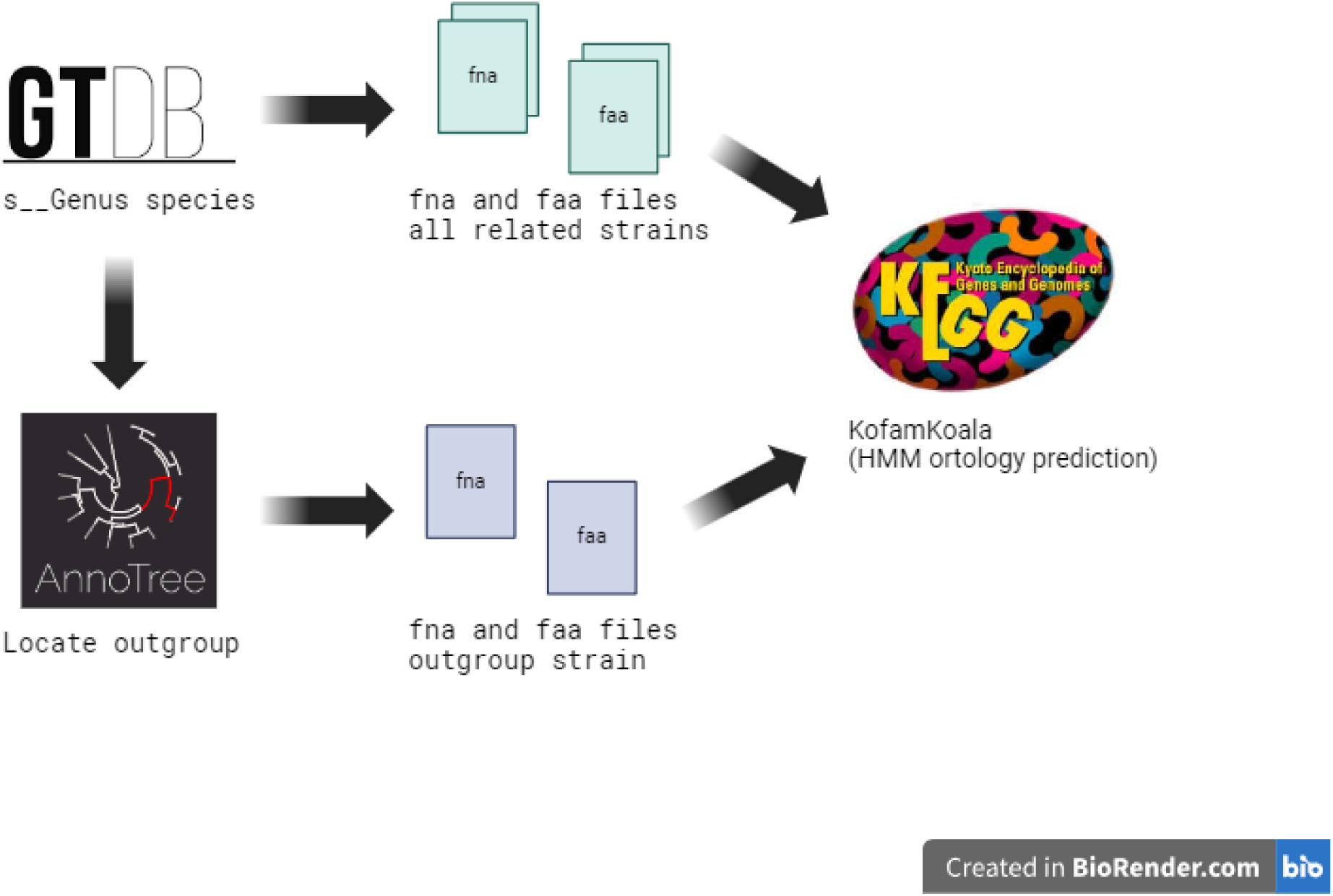
First step of BacNeMu pipeline processing. BacNeMu receives amino acid and nucleotide sequences (fasta files) of all annotated genes from GTDB and automatically identifies an outgroup with AnnoTree. Fasta files can be uploaded manually as well, in this case outgroup will not be automatically located and the AnnoTree part will be ignored. In the case of using the manual loading approach, the outgroup should be placed in input by the user. Then gene orthologs are identified with KofamKoala^28^.

For accurate mutational-spectrum reconstruction, the gene tree used to infer substitutions must reflect the species phylogeny. This condition is satisfied by orthologs — homologous genes whose divergence coincides with speciation events — but is violated by paralogs: genes that arise through intra-genomic duplication and may subsequently diverge in function^29^. Horizontally acquired genes present the same difficulty as paralogs, because they too disrupt the correspondence between the gene tree and the species tree. When non-orthologous sequences are inadvertently included in the alignment alongside orthologs, the resulting phylogeny conflates vertical and non-vertical signals, and substitutions are mapped to incorrect lineages. BacNeMu therefore applies a strict one-to-one (1:1) orthology filter: a gene is retained only if it has exactly one best-matching ortholog in every analysed genome.

To implement this filter, BacNeMu uses the KEGG Orthology database, which organises homologous proteins from diverse species into curated functional clusters, each identified by a unique accession (kegg_id, e.g., K00001)^3^. For each protein in the input set, KofamKoala^28^ (run parameters: --format=mapper-one-line --no-report-unannotated) queries KEGG using hidden Markov model (HMM) profiles and assigns the best-matching cluster. Because many bacterial strains — especially those belonging to the same species — are not yet represented in KEGG Orthology, KofamKoala’s HMM-based search is essential: it can assign a KEGG cluster to a novel protein on the basis of profile similarity, even when the protein lacks a pre-existing database entry.

Once KofamKoala has evaluated the input sequences, BacNeMu performs a stringent filtration step at the level of individual genomes to isolate 1:1 orthologs. First, any gene that KofamKoala cannot confidently assign to a single KEGG cluster (either because it matches multiple clusters with comparable scores, e.g., gene1 matching K0001 and K0003, or because it falls below the homology threshold) is excluded. Second, if multiple distinct genes from the same genome are assigned to the *same* KEGG cluster (e.g., gene2 and gene5 both matching K0002), all such genes are flagged as paralogs and removed for that specific genome. Following these within-genome exclusions, the surviving genes are aggregated into cluster-specific nucleotide FASTA files. If the outgroup strain is missing from a given cluster — either because it naturally lacks the ortholog or because its sequence was filtered out in the previous steps — the entire KEGG cluster is dropped from the analysis, as its gene tree cannot be properly rooted (Fig. 2).

**Figure 2.**
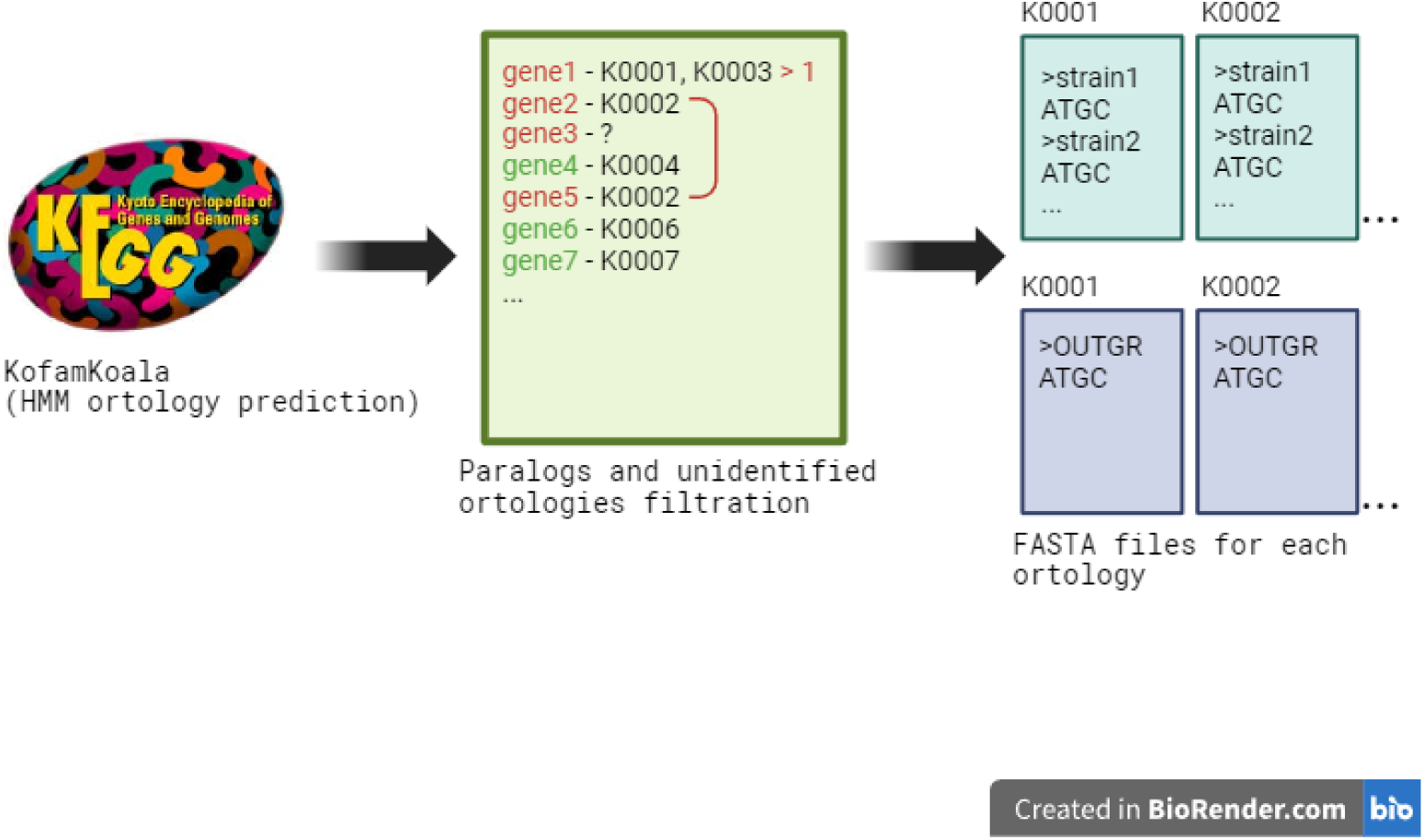
Second step BacNeMu pipeline: gene orthology filtration. BacNeMu filters out paralogs and unidentified orthologies (red colored names), then proceeds the result (green colored names) to separate files named after corresponding KEGG clusters.

For each retained KEGG cluster, the nucleotide sequences of the orthologous genes are collected into a single FASTA file. If the outgroup strain contributes an ortholog to the cluster, its sequence is included in the same file. Sequence duplicates — which carry no additional substitution signal and would artificially compress branch lengths — are removed with seqkit rmdup^30^. Following deduplication, any cluster containing fewer than five sequences (outgroup + four focal strains) is discarded. The remaining alignments are constructed with MAFFT using the --auto parameter, which selects the optimal algorithm based on dataset size and sequence characteristics (Fig. 3).

**Figure 3.**
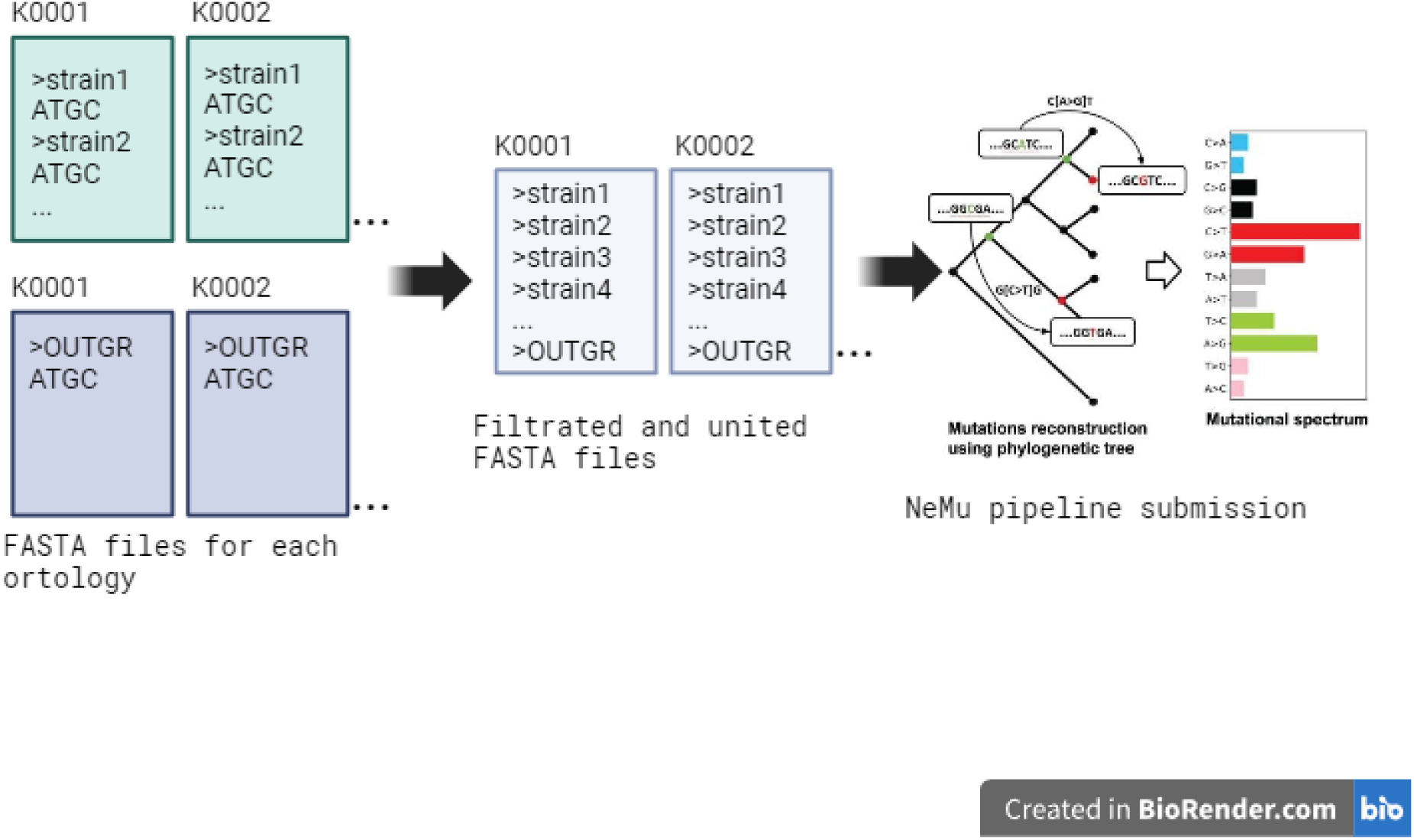
Third step of BacNeMu: sequence abundance filtration and NeMu proceeding part. BacNeMu begins a series of sequence abundance filtration applied for every fasta file resulting from the previous step of the pipeline before proceeding to NeMu. Each file must fit several criteria: have more than 4 not duplicated gene sequences and one outgroup sequence. Fasta files fitting criteria are getting aligned using mafft and proceed to NeMu, resulting in a separate mutational spectrum for each identified orthology.

Each filtered alignment is passed to the NeMu module, which constructs a gene tree using IQ-TREE^25^. Codons are classified by degeneracy, and synonymous substitutions are identified at two-fold, three-fold, and four-fold degenerate sites. By default, only synonymous changes are tallied — reflecting the pipeline’s focus on neutral mutations — but the user may optionally include all substitutions or restrict the analysis to four-fold degenerate sites only, as well as for non-synonymous and all available mutations. Ancestral codon states are reconstructed at every internal node using maximum likelihood, yielding a probability distribution over all possible codons that accounts for branch lengths and the underlying substitution model^4^ (Fig. 3).

For each gene tree, NeMu reconstructs the probability distribution of ancestral nucleotides at every site of every internal node. Each observed substitution is assigned a probability equal to the probability of the ancestral state multiplied by the probability of the descendant state at the corresponding site; substitutions with a probability below a user-defined threshold (0.3 by default) are discarded. Mutations are tallied by substitution type, initially producing a 12-component spectrum that distinguishes the two DNA strands. Because the replicative strand is not known a priori for most bacterial genes, complementary substitutions are merged — yielding a six-component spectrum — by averaging the frequencies of each complementary pair. The pipeline sums the probabilities of all observed mutations of each type and normalises them by the expected mutational opportunities: the sum, over all internal branches and codon sites, of the probabilities of every change permitted under the user-selected codon-type filter (e.g., synonymous only) given the reconstructed ancestral states and the fitted substitution model^4^. This observed-to-expected ratio yields the relative frequency of each substitution type for a single gene.

To produce a species-level spectrum, the gene-specific observed and expected probabilities are summed across all orthologous clusters retained for that species, and the six ratios are recomputed from the pooled counts. Confidence intervals are obtained by bootstrapping over orthological gene clusters: 1,000 resamples of the same size as the original gene set are drawn with replacement, the pooled spectrum is recalculated for each resample, and either the standard error of the mean or percentile-based intervals are reported as the confidence measure. When multiple species are analysed in a single run, BacNeMu also computes an averaged clade-level spectrum. In this case, species-level spectra are treated as independent observations, and the standard error of the mean (SEM) is reported as the confidence measure. All spectra are exported as six-component bar plots, with absolute observed mutation counts annotated on the bars; gene-level spectra are additionally available for users who require finer resolution.

To assess the accuracy of BacNeMu, we compared its species-level spectra against two independent benchmarks: (i) mutation accumulation experiment (MAE) spectra from published studies and (ii) spectra reconstructed by MutTui, a published computational pipeline that infers mutational spectra from population-level SNPs without outgroup rooting or maximum-likelihood ancestral codon reconstruction.

MAE data were obtained for six bacterial species for which experimental mutation-accumulation spectra have been reported: *Escherichia coli* (three non-mutator strains)^31^, *Bacillus subtilis*^32^, *Vibrio cholerae* (two non-mutator strains)^33^, *Aliivibrio fischeri*^33^, *Burkholderia cenocepacia*^34^, and *Mycobacterium smegmatis*^35^. In MAE studies, the mutation rate for each substitution type is typically reported as the number of mutations per nucleotide per generation. For consistency with the relative-frequency format of BacNeMu and MutTui outputs, these per-nucleotide-per-generation rates were taken from the supplementary materials of the cited publications and converted to relative frequencies by dividing each substitution-specific rate by the sum of all six rates, yielding a six-component vector that sums to unity. For species represented by multiple MAE strains, each strain was treated as a separate observation and compared individually with the corresponding species-level BacNeMu spectrum.

MutTui spectra were reconstructed from the same set of strain identifiers used for BacNeMu, ensuring a matched comparison. MutTui assigns the direction of each SNP along a precomputed phylogeny using TreeTime^36^ and rescales the resulting mutation counts by the genomic nucleotide composition of the reference genome. In contrast, BacNeMu employs outgroup rooting and maximum-likelihood reconstruction of ancestral codons, and normalises mutation counts as an observed-to-expected ratio (see above). By default, BacNeMu tallies only synonymous substitutions, whereas MutTui reports all substitutions irrespective of codon context; accordingly, we ran BacNeMu in two configurations — synonymous-only and all-mutations — when computing pairwise similarities (see Results Fig. 4).

**Figure 4.**
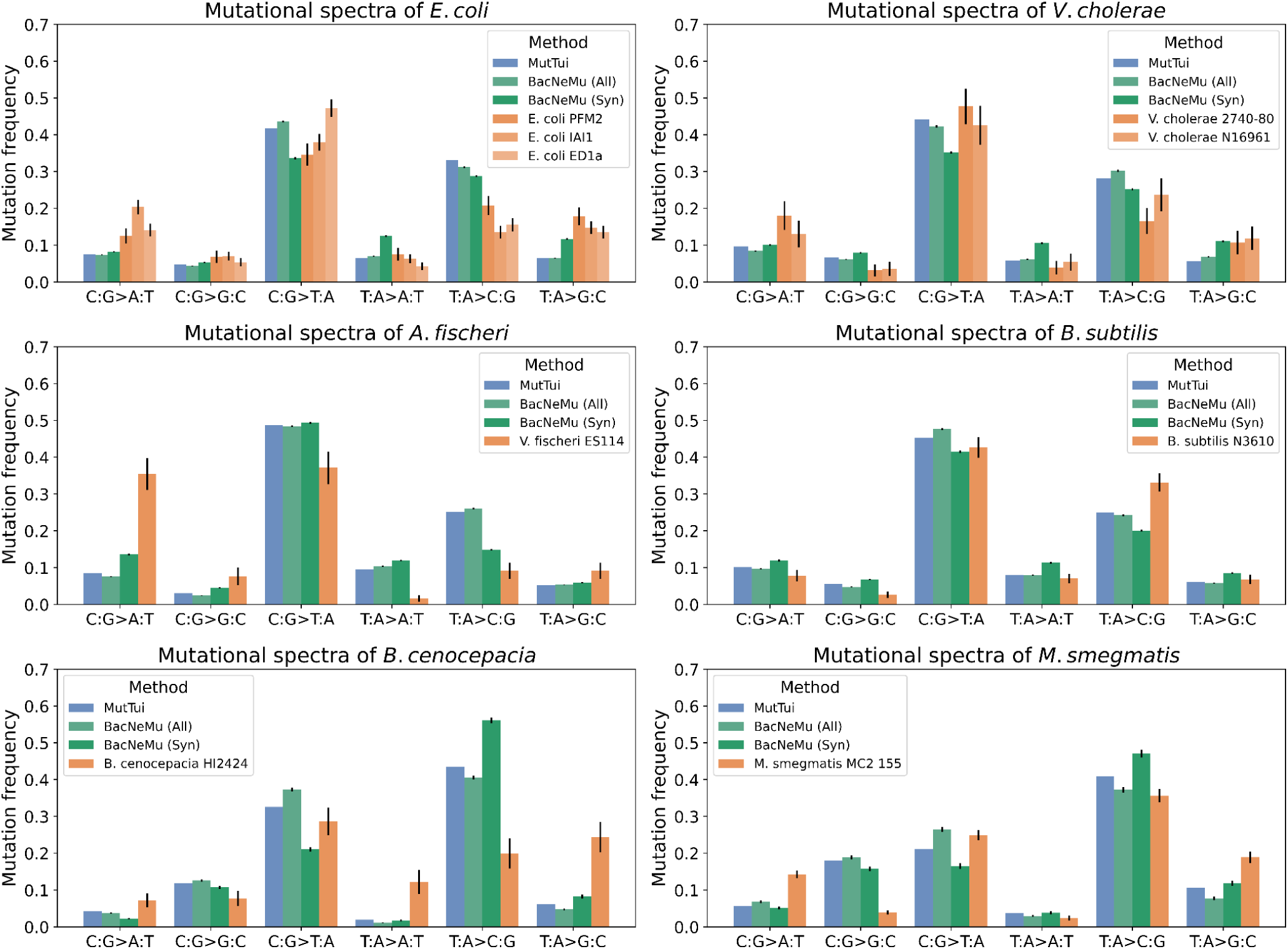
Comparison of mutational spectra reconstruction methods shows existing affinity between MutTui, BacNeMu (All and Syn) and mutation accumulation experiment approaches. Blue - MutTui with GC-content normalization. Green - BacNeMu pipeline reconstruction. Orange - mutation accumulation experiment strain.

Pairwise agreement between spectra was quantified using cosine similarity, which captures the similarity of the relative proportions of the six substitution types independently of the absolute mutation counts. The metric ranges from 0 (orthogonal vectors) to 1 (identical directionality). A matrix of pairwise cosine similarities among all three approaches — BacNeMu, MutTui, and MAE — was assembled for each species and is presented in Results (Fig. 5).

**Figure 5.**
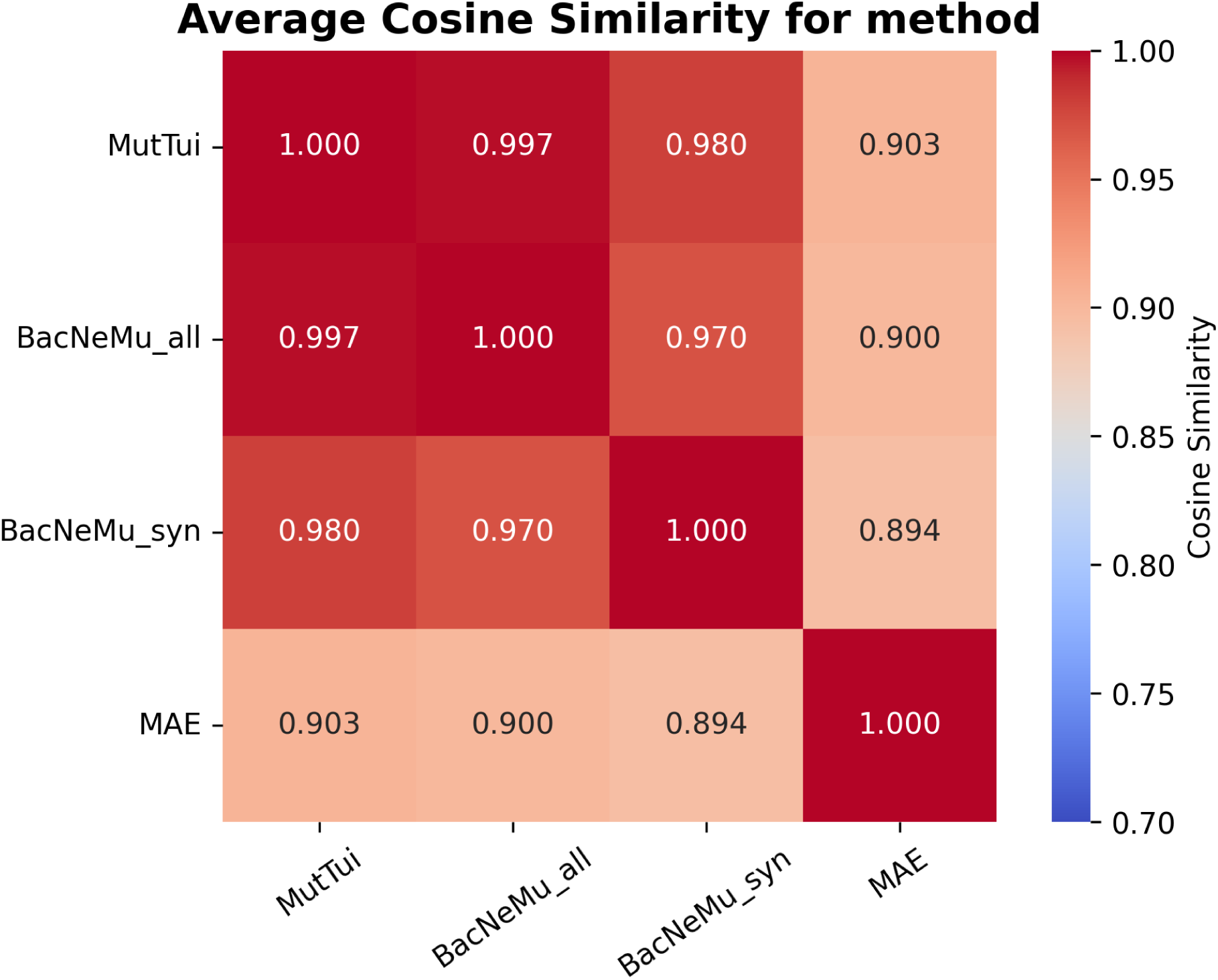
Matrix of averaged cosine similarities of different methods of mutational spectra reconstruction. The shown values are averages across all corresponding species mutational spectra comparisons (see Supplementary materials).

## Results

### Pipeline overview

The BacNeMu pipeline (Fig. 1–3) reconstructs six-component neutral mutational spectra for bacteria from a user-provided set of bacterial taxa (one or several). The pipeline (i) accepts the bacterial taxa of interest; (ii) collects all annotated protein-coding gene sequences and their corresponding protein sequences for the given taxa; (iii) automatically identifies an outgroup strain as the closest reference genome that does not belong to the focal species (Fig. 1); (iv) retains only one-to-one orthologous gene clusters; (v) keeps only those clusters represented by at least four non-outgroup strains; (vi) generates codon-based alignments for each orthologous cluster (Fig. 2); (vii) reconstructs a mutation spectrum for each orthologous cluster using the previously developed NeMu framework — building a gene tree with IQ-TREE, inferring ancestral states at internal nodes using the outgroup, and estimating the frequency of each mutation type from synonymous substitutions; (viii) aggregates the gene-specific mutation spectra into a species-specific mutation spectrum; (ix) provides comprehensive, configurable output for the six possible dinucleotide transitions, i.e. the six-component mutational spectrum.

The pipeline supports both automatic and manual modes, offers numerous adjustable parameters, and allows output at different levels of detail. Run time ranges from 1 to 8 hours depending on the number of input taxa (minimal input of 4 strains — approximately 1 hour; maximum recommended input of 100 strains — approximately 8 hours) and on the computational configuration. Further details are provided in Materials and Methods.

### Pipeline validation

BacNeMu was validated by comparing its reconstructed spectra against two independent benchmarks: published mutation accumulation experiment (MAE) data and spectra reconstructed by MutTui^26^, an alternative computational pipeline. MAE data were obtained for six bacterial species — *Escherichia coli* (three non-mutator strains)^31^, *Bacillus subtilis*^32^, *Vibrio cholerae* (two non-mutator strains)^33^, *Aliivibrio fischeri*^33^, *Burkholderia cenocepacia*^34^, and *Mycobacterium smegmatis*^35^ — yielding nine individual strain-level spectra, which were not averaged across strains of the same species. BacNeMu spectra were reconstructed with default parameters and normalised so that the six substitution frequencies summed to unity; confidence intervals were obtained by bootstrap resampling (1,000 iterations) as described in Materials and Methods. MutTui spectra were generated from the same set of strain identifiers; methodological details of the comparison are provided in Materials and Methods.

The resulting mutational spectra are shown in Figure 4. Across all six species, the spectra produced by MutTui and by BacNeMu in all-mutations mode were nearly identical, while BacNeMu in synonymous-only mode exhibited small deviations that are expected given its restriction to neutrally evolving sites, yet preserved the overall spectral shape. The experimentally determined MAE spectra displayed greater strain-to-strain variability — most pronounced in *B. cenocepacia* and *M. smegmatis* — though the dominant substitution types remained consistent with those recovered computationally. This variability is expected given the fundamental difference in timescale between the two approaches: MAE captures mutations arising over hundreds of generations in a single clonal lineage under laboratory conditions, whereas phylogenetic methods integrate fixed substitutions accumulated over evolutionary time across multiple diverse strains. Despite these differences, the overall concordance between the computational and experimental spectra suggests that the phylogenetic approach yields a reliable approximation of the underlying neutral mutational spectrum (see Discussion). Pairwise agreement between methods was quantified using cosine similarity (Fig. 5).

The closest correspondence was observed between MutTui and BacNeMu in all-mutations mode, with an average cosine similarity of 0.997 across species (Fig. 5). Similarities between the computational methods and the MAE spectra were also high and comparable to each other, indicating that BacNeMu recovers mutational frequencies with an accuracy comparable to that of established bioinformatic tools and produces spectra broadly consistent with experimental MAE data.

### Case studies

#### Obligate Aerobe vs Obligate Anaerobe

Aerobic organisms are expected to experience more frequent oxidative damage than anaerobic organisms, and, due to oxidatively induced G>T (G:C>T:A) transversions^14,37,38^, aerobiosis has traditionally been thought to decrease GC content. However, multiple studies^14,39–46^ have shown that, contrary to expectations, aerobic prokaryotes actually have higher GC contents than anaerobic prokaryotes. After many years of this paradox existed, it has been hypothesized that, in prokaryotes, oxidative damage may produce additional mutational signatures beyond or instead of G>T (G:C>T:A) transversions, shifting aerobic genomes towards higher GC content^15^. Notably, one such oxidative damage candidate signature, A>G (A:T>G:C), has been described in several recent studies of mitochondrial DNA^4,5,47^, which shares a prokaryotic origin as well as in yeast single-strand DNA^6^. Here, to perform a pilot test of potentially increased A>G substitutions in aerobes versus anaerobes, which may help resolve the long-standing paradox between oxidative damage and genomic GC content, we demonstrate the functionality of BacNeMu to reconstruct the mutational spectra.

To demonstrate the functionality of the developed pipeline, we tested it on bacterial clades with well-characterised oxygen requirements, spanning obligate aerobes and obligate anaerobes. Consistent with a previously published report demonstrating a significant predominance of guanine and cytosine nucleotides in obligate aerobes^15^, the mutation spectra of obligate aerobes reconstructed using the BacNeMu pipeline exhibited a predominance of A>G (T>C) transitions (Fig. 6). To begin with, we reconstructed the mutation spectra of three bacterial species: *Clostridium cagae* - shown in the study as gram-positive strictly obligate anaerobe^48^; *Mycobacterium tuberculosis* - parasite, commonly classified as aerobe, but with evidence of anaerobic strains existence^49^. *Cupriavidus basilensis* - strictly aerobic bacterium, confirmed to perish rapidly in anaerobic environments^50^. The resulting mutational frequencies (Fig. 6) showed that obligate aerobe *Cupriavidus basilensis* has higher T:A>C:G than *Clostridium cagae*, as expected.

**Figure 6.**
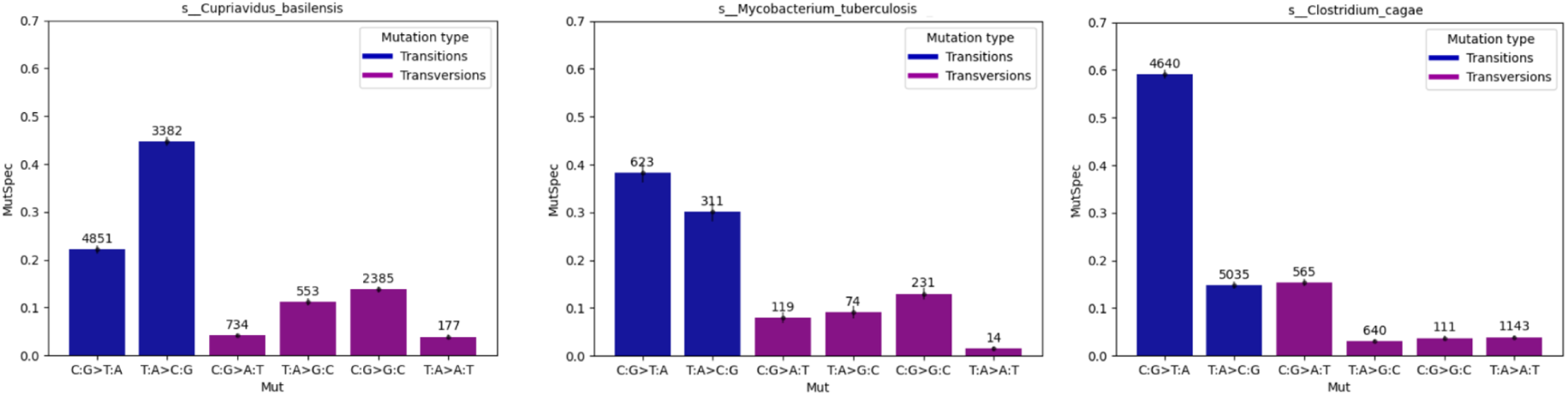
Mutational spectra of 3 randomly selected species based on their aerobic status confirmed with corresponding publication (obligate aerobic, parasitic semi-aerobic and obligate anaerobic) reconstructed using BacNeMu pipeline with only synonymous mutations.

The results obtained for individual species are consistent with the initial hypothesis. However, for a more in-depth analysis, the utilization of the JGI GOLD database^51^ as a large-scale, validated source of biological determinants enables broadening of the input data of obligate aerobic and obligate anaerobic species. We reconstructed all available obligate aerobic and obligate anaerobic species (264 obligate anaerobes and 132 obligate aerobes) from the JGI GOLD database, and then calculated the average mutation spectra from the species-level data for each clade. As a result, mutational spectra were calculated for 28 obligate anaerobes and 16 obligate aerobes. This shrinkage of species number arises from the insufficient volume of annotated bacterial genomes currently available in the GTDB database. However, the estimate can be considered reliable, because with the exception of two clades, all remaining species are widely distributed across the phylogenetic tree, which completely rules out the influence of phylogenetic inertia in the analysis. In future studies we are going to expand the number of available species with multiple databases (see Discussion). Standard error whiskers were used to show statistical significance. The average mutational spectrum of the obligate aerobes shows a higher frequency of T:A>C:G and lower frequency of C:G>T:A transversions compared to the obligate anaerobes (Fig. 7A). In order to have a closer look into mutational spectra distribution across taxa, bacterial tree of life AnnoTree was used to show all species reconstructed in this comparative analysis (Fig. 7B). The tree shows that some obligate aerobic and obligate anaerobic species show opposite mutational spectrum from expected, which may occur due to specifics of repair system evolution or environment conditions of some species and requires deeper analysis.

**Figure 7.**
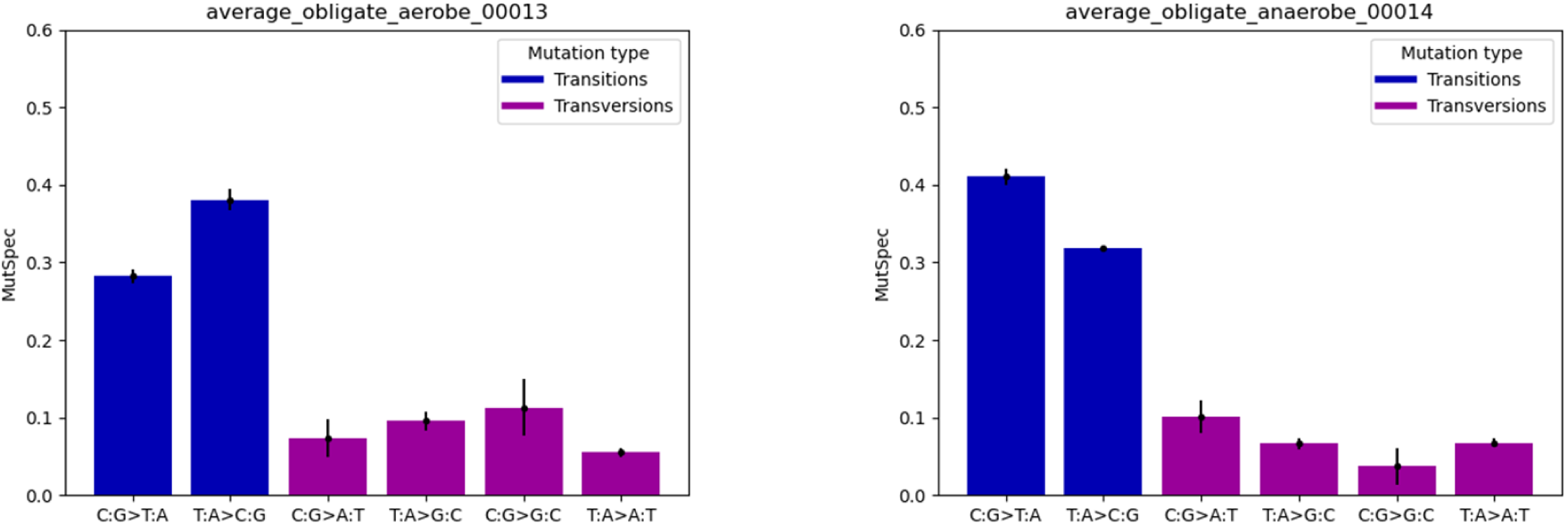
A. Average mutational spectrum of obligate aerobic and obligate anaerobic species from JGI GOLD and AnnoTree.

**Figure 7.**
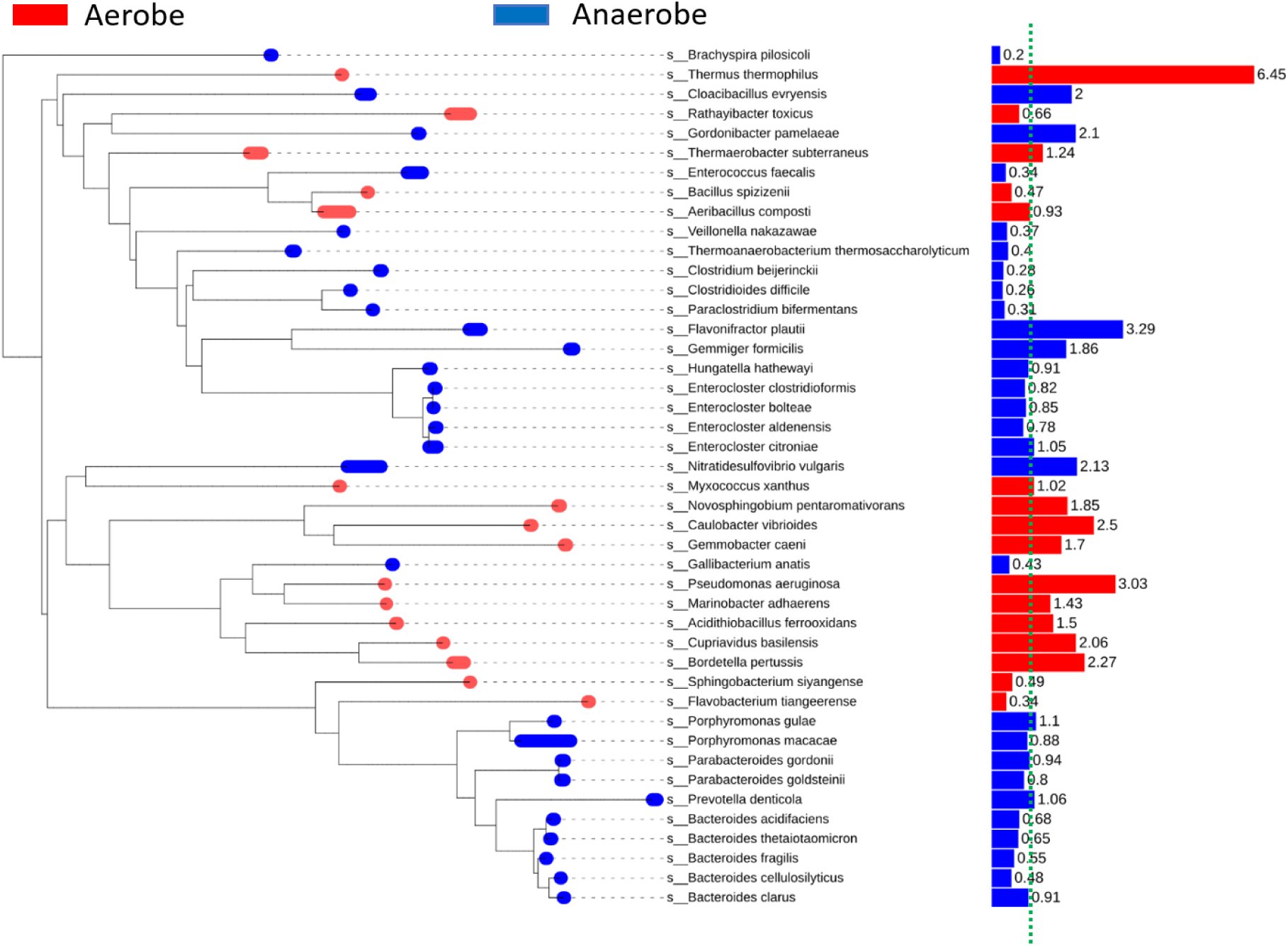
B. Obligate aerobes and anaerobes phylogenetic tree visualization. Red branch dot - JGI GOLD obligate aerobe annotation. Blue branch dot - JGI GOLD obligate anaerobe annotation. The colour code for barplots is the same. We expected A:T>G:C/G:C>A:T rate to be more than 1 (green line) for obligate aerobes (red barplots) and less than 1 for obligate anaerobic species (blue barplots), which is not always the case, however true on average (Fig. 7A)

As evident from the phylogenetic tree shown above, certain species display mutation spectra that deviate markedly from the average characteristic of their oxygen status. To confirm the robustness of these results, two-tailed and one-tailed U-tests were performed, revealing significantly higher A:T>G:C/G:C>A:T rates in obligate aerobes than in obligate anaerobes (two-tailed U-test p-value: 0.019 < 0.05; one-tailed U-test p-value: 0.00958 < 0.01). Furthermore, a phylogenetic generalized least squares (PGLS) analysis, conducted to correct for the effect of closely related obligate anaerobe clades, yielded moderate significance (p-value = 0.17). The mutational frequencies are also shown in Table S1, which demonstrates several additional significant differences among transitions between obligate aerobes and obligate anaerobes.

#### Thermophile vs psychrophile

In addition to oxygen requirements, the influence of the other ecological factors can also be investigated using the BacNeMu pipeline. A subject of active debate in the scientific literature is the elevated GC content observed in thermophiles compared to psychrophiles, which may indicate an influence of temperature on bacterial DNA stability^16^. No distinct pattern has been established for psychrophiles, and within the framework of this study, it cannot be expected that C:G>T:A transitions will be as elevated in psychrophiles as was observed in obligate anaerobes. Furthermore, the identification of psychrophiles poses an additional challenge, as their annotation is performed without a standardized protocol, which may lead to spurious results in comparative analyses^52^. For this experiment, three random species were selected: one designated as a hyperthermophile, another as a hyperpsychrophile, and the third which grows in moderate temperature interval (4-37°C) and serving as a control for comparison.

Three bacterial species were selected for a quick experiment: *Thermotoga maritima* - hyperthermophilic bacteria, which is known to be able to grow in high temperatures - 70-90°C^7^; *Psychrobacter sanguinis* - cold-resistant parasite bacteria^9^; *Clostridium algidicarnis* - psychrophile, bacteria isolated from meat after vacuum freezing^8^. Mutational spectra show prevalence of A>G (T>C) mutation in thermophilic bacteria mutspec compared to others (fig. 8).

**Figure 8.**
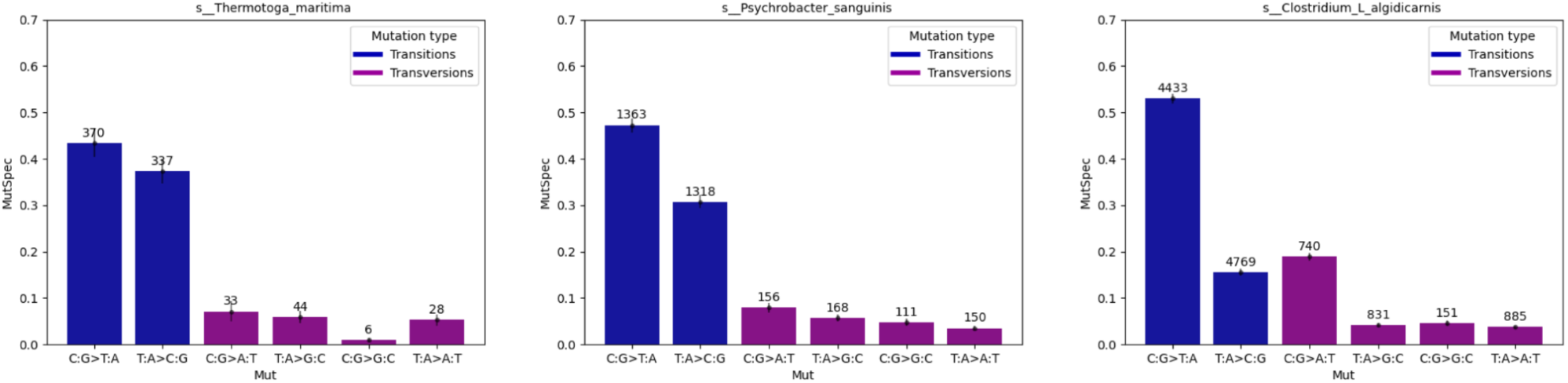
Mutational spectrum of 3 randomly selected species based on their temperature status (*Thermotoga maritima* - hyperthermophile, *Clostridium algidicarnis* - hyperpsychrophile and *Psychrobacter sanguinis* - parasitic psychrophile) reconstructed using BacNeMu pipeline.

The mutation spectrum of the hyperthermophile *Thermotoga maritima* reveals a predominance of C:G>T:A transitions over T:A>C:G. In contrast, the hyperpsychrophile *Clostridium algidicarnis* exhibits a strong predominance of C:G>T:A over T:A>C:G, along with a notably high frequency of C:G>A:T transversions; the substantial number of mutations supports the reliability of this result. The control species *Psychrobacter sanguinis*, despite being designated as psychrophilic, also displays a marked predominance of C:G>T:A, yet its T:A>C:G frequency is higher than that observed in the hyperpsychrophile *C. algidicarnis*. Collectively, this study demonstrates that the BacNeMu pipeline can yield biologically meaningful results from relatively small bacterial samples; nevertheless, drawing definitive conclusions regarding the influence of temperature on mutagenesis will require a larger-scale investigation that has not been yet conducted. This work may ultimately help address the basis of GC-rich genomes in thermophiles, while a key challenge for existing phylogenetic algorithms reconstructing mutation spectra remains the need to disentangle the effects of natural selection on nucleotide composition from those of neutral mutagenesis.

## Discussion

In this work, we introduced BacNeMu — a phylogenetically informed pipeline for reconstructing neutral mutational spectra of bacteria from publicly available genomic databases. Our validation against both mutation accumulation experiments and the independent computational method MutTui^26^ demonstrates that BacNeMu recovers biologically faithful six-component spectra while reducing the time required from multiple months or years (in the case of MAE) to a matter of hours. Applied to obligate aerobes and anaerobes, the pipeline successfully detected the expected excess of A:T>G:C transitions in aerobic species. This finding is consistent with the hypothesis that oxidative damage biases the mutational process toward A:T>G:C substitutions. Rather, it suggests that mutational bias acts in the same direction as — and may complement — selection for GC-rich genomes in oxygen-exposed prokaryotes^18^. The thermophile–psychrophile pilot, although limited in scale, further illustrates the pipeline’s utility for probing the relationship between ecological variables — such as growth temperature — and the mutational process. Taken together, these results establish BacNeMu as a practical, high-throughput tool for comparative mutagenesis studies in bacteria, capable of generating testable hypotheses about how environmental pressures shape the mutational landscape.

Several limitations of the current implementation point to directions for future work. First, BacNeMu currently collapses the full 12-component strand-specific spectrum into a six-component spectrum by averaging complementary substitution pairs. This simplification is necessitated by the absence of reliable replicative-strand annotation for most bacterial genes, yet it discards potentially informative asymmetry between the leading and lagging strands; extending the pipeline to infer strand orientation — for example, by incorporating origin-of-replication data or gene orientation biases — would enable 12-component analyses. Second, our reliance on GTDB as the primary source of annotated genomes, while ensuring taxonomic consistency, substantially limited the number of species available for the aerobic–anaerobic comparison: of the 396 candidate species identified from JGI GOLD, only 44 could be processed due to lack of available annotated genomes. Integrating additional genome repositories — such as GenBank — would expand taxonomic coverage and increase the number of retained orthologous clusters per species, improving statistical power. Furthermore, the pipeline currently depends on JGI GOLD for ecological annotations (e.g., oxygen tolerance), which covers only a fraction of bacterial diversity. Incorporating the BacDive^53^ database — a curated resource of strain-level metabolic and ecological traits — would substantially increase the number of species with reliable phenotype assignments and enable more nuanced analyses, such as distinguishing obligate from facultative anaerobes. Beyond these methodological extensions, several biological questions become accessible with BacNeMu and warrant investigation: (i) whether bacteria inhabiting high-UV environments exhibit elevated C:G>T:A transitions^54^, as predicted by UV-induced pyrimidine dimer formation^55^; (ii) whether intracellular parasites with reduced genomes display mutational spectra that differ systematically from those of free-living relatives, potentially reflecting relaxed selection on DNA repair pathways^56–60^; and (iii) whether the mutational spectra of antibiotic-resistant clinical isolates differ from those of susceptible strains, which could shed light on the role of hypermutation in the emergence of resistance^61^.

---

**Table S1.** Pairwise comparisons of individual mutational spectrum components between obligate aerobes and obligate anaerobes.

| Mutation | Median (Aerobe)<br>(Q1 = Q3 interval) | Median<br>(Anaerobe)<br>(Q1 = Q3 interval) | two-tailed U-test<br>p-value |
| --- | --- | --- | --- |
| C:G>T:A | 0.26 | 0.405 | 0.001 |
| T:A>C:G | 0.365 | 0.305 | 0.140 |
| C:G>A:T | 0.075 | 0.09 | 0.098 |
| T:A>G:C | 0.09 | 0.07 | 0.001 |
| C:G>G:C | 0.11 | 0.035 | 3.46e-06 |
| T:A>A:T | 0.03 | 0.07 | 0.098 |

